# Mass spectrometry imaging reveals early metabolic priming of cell lineage in differentiating human induced pluripotent stem cells

**DOI:** 10.1101/2022.08.16.504020

**Authors:** Arina A. Nikitina, Alexandria Van Grouw, Tanya Roysam, Danning Huang, Facundo M. Fernández, Melissa L. Kemp

## Abstract

Induced pluripotent stem cells (iPSCs) hold great promise in regenerative medicine; however, few algorithms of quality control at the earliest stages of differentiation have been established. Despite lipids having known functions in cell signaling, their role in pluripotency maintenance and lineage specification is underexplored. We investigated changes in iPSC lipid profiles during initial loss of pluripotency over the course of spontaneous differentiation using co-registration of confocal microscopy and matrix-assisted laser desorption/ionization (MALDI) mass spectrometry imaging. We identified lipids that are highly informative of the temporal stage of the differentiation and can reveal lineage bifurcation occurring metabolically. Several phosphatidylinositol species emerged from machine learning analysis as early metabolic markers of pluripotency loss, preceding changes in Oct4. Manipulation of phospholipids via PI 3-kinase inhibition during differentiation manifested in spatial reorganization of the colony and elevated expression of NCAM-1. In addition, continuous inhibition of phosphatidylethanolamine N-methyltransferase during differentiation resulted in increased pluripotency maintenance. Our machine learning analysis highlights the predictive power of metabolic metrics for evaluating lineage specification in the initial stages of spontaneous iPSC differentiation.

## Introduction

Induced Pluripotent Stem Cells (iPSCs) can be reprogrammed from a patient’s own adult cells [1] and differentiated into any cell type with many potential clinical uses ([2], [3], [4], [5]). Numerous *in vitro* studies have explored various differentiation protocols, resulting in tissues of interest (e.g. [6], [7], [8], [9]). Spontaneous, or undirected, differentiation allows production of all three germ lineages and can be used as a model of initial loss of pluripotency that is applicable to a wide range of protocols. Human iPSC colonies are disordered, unlike embryos, yet take on a degree of self-assembly and organization over time; however, the mechanisms of cellular reprogramming and colony self-organization are still understudied. When iPSCs are used for regenerative medicine, quality control and a thorough understanding of the occurring intracellular processes are essential to prevent carcinogenesis and contamination by unwanted cells.

In a cell manufacturing setting, typical quality control includes initial confirmation of cellular pluripotency by confirming sufficient Oct4 expression in the colony sample [10]. After a differentiation protocol is completed, quality control can include quantifying expression levels of phenotype marker genes by flow cytometry as well as tissue functional tests (e.g. contractility in cardiomyocytes, production of collagen in fibroblasts, etc.). More extensive quality control of a finalized clinical treatment can include whole genome sequencing and whole exome sequencing [5].

Most of the described approaches are destructive, with only several known glycoprotein surface markers allowing real-time quality control. To date, quality control has rarely been performed by assessing cellular lipids, and the importance of lipidome remodeling has generally been overlooked. Recently, expression of plasmalogens and sphingomyelins was shown to increase during the process of iPSC differentiation into vascular endothelial cells [11], suggesting that phospholipid metabolism plays an important role. In addition to their well-known contribution to structure in membranes, polyunsaturated phospholipids are precursors of important signaling molecules [12], and lipid supplementation was previously shown to influence general iPSC phenotype [13]. Here, we focus on glycerophospholipids such as phosphatidic acids (PA), phosphatidylethanolamines (PE), phosphatidylcholines (PC), phosphatidylserines (PS), and phosphatidylinositols (PI). In addition to functioning as negatively charged building blocks of membranes, phosphatidylinositols and related phosphates have crucial roles in the interfacial binding of proteins and in the regulation of protein activity at the cell interface. A well-known example is the Akt/PKB signaling pathway, which is activated by PI 3-kinase phosphorylation of phosphatidylinositols, followed by the recruitment of Akt to the membrane due to the interaction with the resulting phosphoinositide docking sites. Activated Akt then controls many key cellular functions, including differentiation, proliferation, metabolism and apoptosis.

In this work, we assess changes in phospholipid abundances in iPSCs over the course of the spontaneous differentiation protocol as well as their spatial distribution inside a colony using both high and ultrahigh resolution matrix-assisted laser desorption/ionization (MALDI) mass spectrometry (MS) imaging co-registered with confocal microscopy. MALDI MS imaging has been successfully used before to show that the distribution of phosphatidylcholines differs between the differentiated and undifferentiated parts of iPSC colonies [14]. We developed a variety of machine learning models that indicate dynamic and spatial trends at the single-cell lipidome level robustly predict pluripotency loss; furthermore, these modeling tools capture bifurcation in lineage specification between SSEA1+ and NCAM1+ phenotypes.

## Results

### Dynamic changes in phospholipid abundance reveal early metabolic markers of differentiation

To determine the dynamic changes in lipids during loss of pluripotency in iPSCs, we analyzed iPSC colony samples undergoing 0 to 7 days of the spontaneous differentiation protocol using a multi-modal image co-registration pipeline merging immunofluorescence and MALDI MS data to yield high-dimensional metabolic imaging with near-single cell resolution (Methods S2, S5). Briefly, colonies were stained with 3 fluorophore-conjugated antibodies: TRA-1-81 pluripotency marker, SSEA-1 loss of pluripotency marker, and NCAM-1 neural lineage marker; the nuclei were labeled with Hoechst stain. For each of 8 consecutive days of spontaneous differentiation, confocal microscopy and MS images of the same ROI were acquired (Fig. 1a). Next, the two imaging modalities were aligned for each of 8 pairs of colony images, the nuclei in each confocal image were segmented and their contours overlaid on the MS images: the average signal for each selected *m/z* value was then calculated for each nucleus. This protocol yielded 8 datasets with about 10 thousand cells each and 70 *m/z* peak-picked features each. To assess temporal changes in phospholipid abundance occurring during pluripotency loss, we calculated the average signal per day of differentiation for each of the *m/z* features. Nine representative trajectories of interest are shown in Fig. 1b: the abundances of *m/z* 528.3, 722.5 and 748.5 exhibited stable growth with differentiation time, while the abundances of *m/z* 742.55, 778.53, 861.5 and 863.5 showed some initial growth but declined for the remaining differentiation time. The abundance of *m/z* 885.6 was stable for the first 4 days after which it exhibited rapid growth, making it anticorrelated (R = -0.85) with Oct4 expression levels measured *via* flow cytometry (Fig. 1d). The abundance of *m/z* 940.6 rapidly decreased to near-zero values in the first 4 days of differentiation, preceding the reduction in Oct4 expression, suggesting that this species could be an early marker of pluripotency loss. To emphasize the scale of phospholipid abundance changes during spontaneous differentiation, we trained a decision tree classifier (Fig. 1c) using cell-by-cell lipid abundances as features and the day of differentiation as a class label. We used a biological replica of the same experiment as a validation dataset, which yielded 67% validation accuracy when classified into 8 days of spontaneous differentiation. However, the structure of the fitted tree suggested three main branches: days 0-2, 3-5 and 6-7. We labeled these branches as “pluripotent”, “undergoing differentiation” and “differentiated”. With these 3 classes the simplified decision tree yielded 87% validation accuracy in prediction of iPSC state from 7 metabolic features.

**Figure 1.**
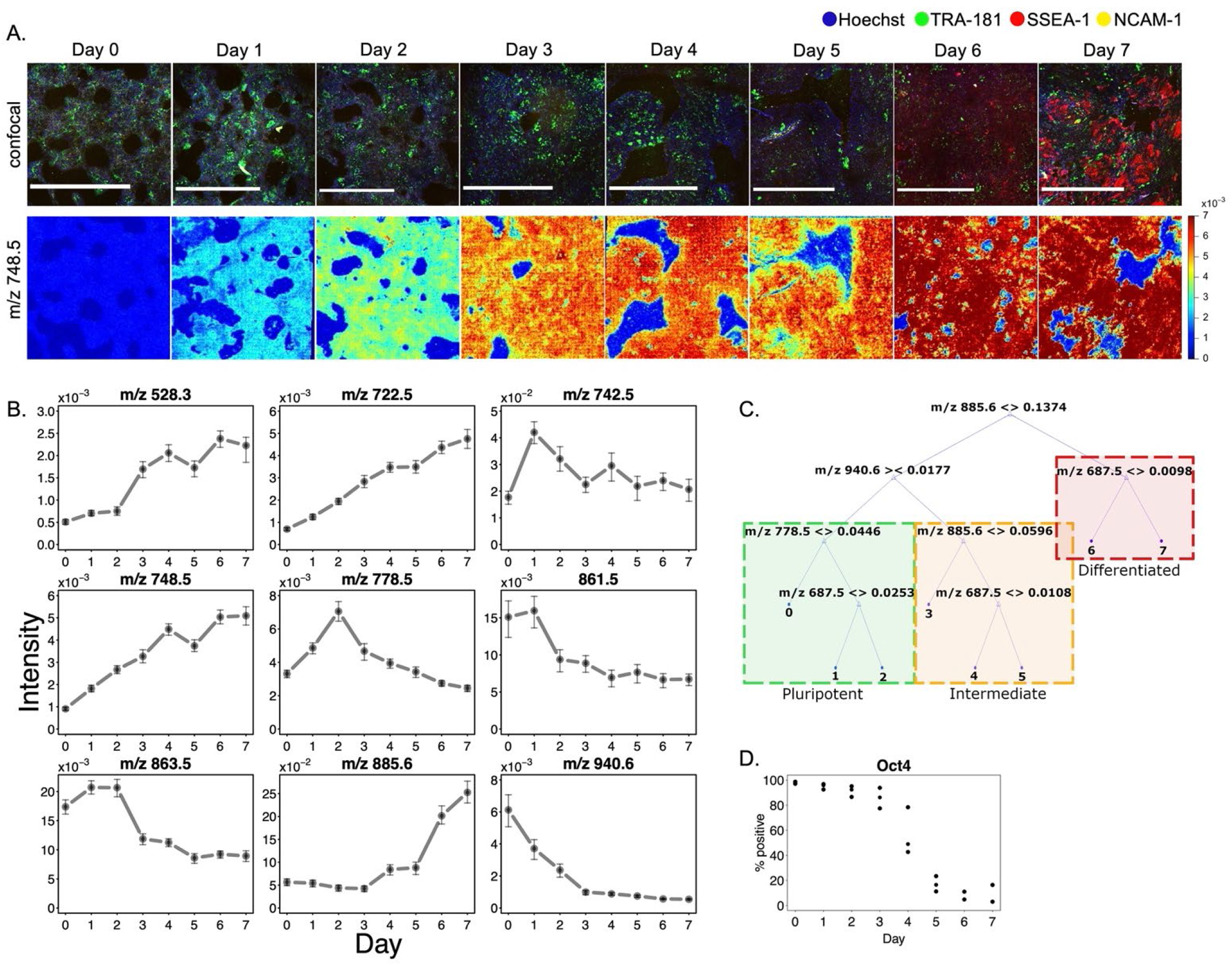
Degree of spontaneous differentiation of iPSC colonies can be predicted through a subset of metabolic features. A. Top row - confocal images of iPSC colonies undergoing differentiation for 7 days, bottom row – corresponding MALDI TOF ion images for *m/z* 748.5. Scalebars are 1 mm. B. Nine examples of temporal changes in mean phospholipid abundance during the differentiation. Error bars show 25 and 75 percentiles. C. Decision tree trained to predict the day of differentiation based on phospholipid abundance with validation accuracy of 67% for classification into 8 days and 87% for classification into 3 major classes: pluripotent, undergoing differentiation and differentiated. For sample sizes refer to Table 1. D. Changes in percent of Oct4 positive cells over 7 days of spontaneous differentiation measured by flowcytometry. Each day had three biological replicates.

### PLS discriminant analysis reveals spatial correlation of phospholipid abundance and pluripotency markers

To associate pluripotency status of an iPSC with its metabolic signature, we analyzed the spatial correlation of *m/z* features with fluorescent pluripotency labels in the imaged colonies. We selected day 6 of spontaneous differentiation for analysis because the cell colony was exhibiting significant expression of both TRA-181 and SSEA-1 pluripotency markers. None of the days showed expression of NCAM-1. Cells in the training sample (Fig. 2a, left side) were labelled as TRA-181 positive or SSEA-1 positive based on k-means clustering (K=2) of the respective fluorescent intensities. We used an experimental replicate of day 6 as the validation dataset (Fig. 2a, right side). We trained a PLS-DA classifier (Fig. 2c) and, after variable trimming, the validation accuracy was 90%. The predicted cell labels are plotted in Fig. 2a alongside the original confocal images. The spatial distribution of *m/z* 742.5 abundance closely correlated with TRA-181 expression (Fig. 2b, left side), in agreement with its decline with differentiation time shown in Fig. 1a. The spatial distribution of *m/z* 885.6 abundance (Fig. 2b, right side) reflected an anticorrelation to SSEA-1 expression in the colony center changing to correlation closer to the edge.

**Figure 2.**
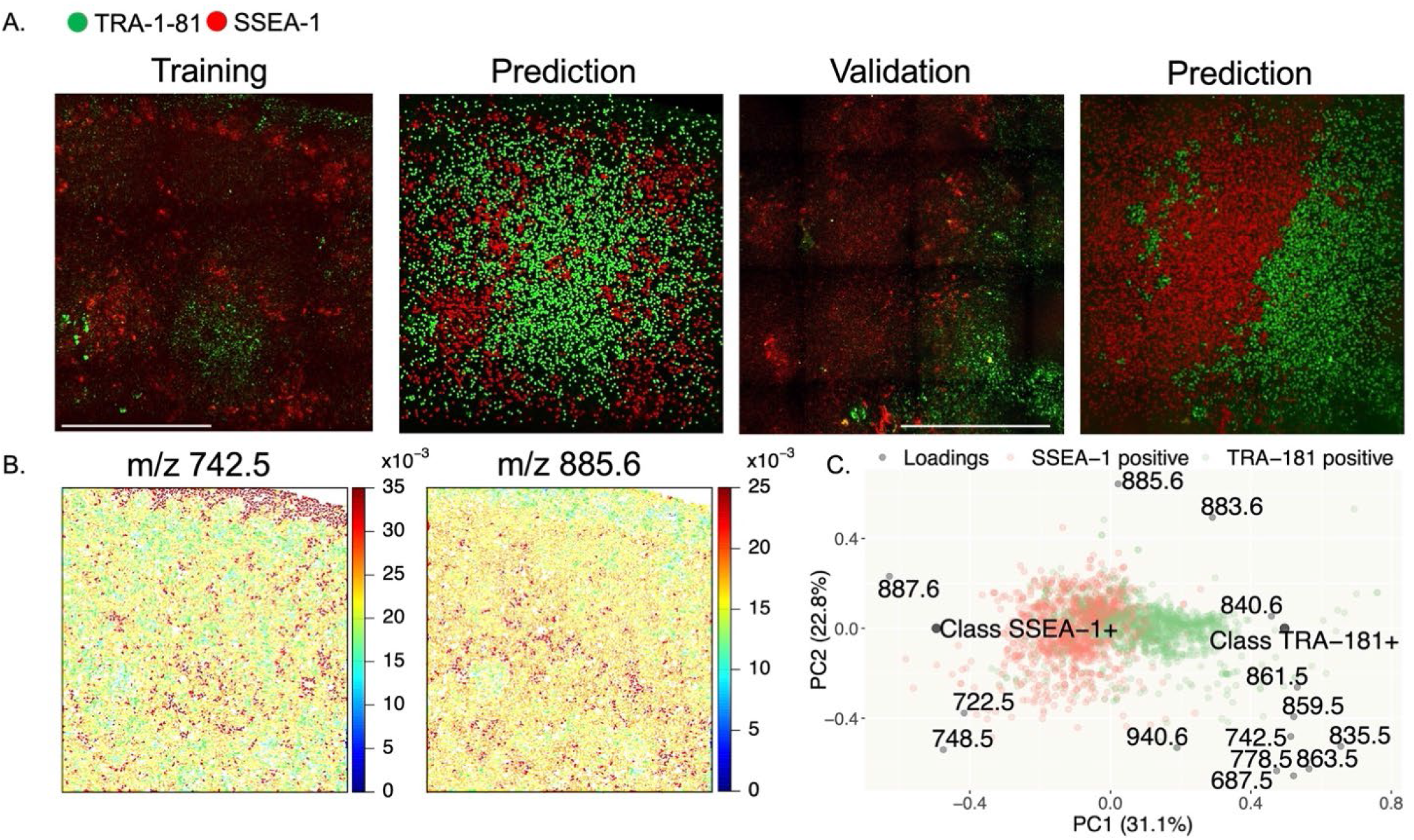
PLS discriminant analysis reveals spatial correlation between phospholipid abundances and pluripotency markers. A. Training and validation confocal images of day 6 of spontaneous differentiation and their predicted pluripotency labels. Green color labels pluripotent cells, red color labels differentiated cells. Scalebars are 1 mm. B. Examples of *m/z* features correlating with cell pluripotency labels. C. Biplot of PLS-DA model used to discriminate between SSEA-1+ and TRA-181+ populations based on cells’ phospholipid abundance with 90% validation accuracy. For sample sizes refer to Table 1.

### Inhibition of phosphatidylethanolamine N-methyltransferase prolonged pluripotency during spontaneous differentiation

We annotated as many detected lipids as possible through MS/MS experiments and accurate mass measurements (Table 2) to relate the metabolic features with biological function. Several phospholipids with abundance changes associated with the differentiation process were annotated as phosphatidylethanolamines (PEs). With previous studies suggesting that phosphatidylcholines (PCs) are involved in differentiation [14] we disrupted the PE to PC conversion pathway by inhibiting PEMT by addition of 50 *µ*M of 3-Deazaadenosine (DZA) to the differentiation media throughout all 7 days of differentiation. We observed via flow cytometry (Fig. 3) that continuous DZA exposure prevents Oct4 expression loss with differentiation. To reveal changes in phospholipid abundances following this perturbation, we grew an additional 8 iPSC colony samples, one for each day of spontaneous differentiation with constant DZA exposure. Since we did not observe any changes in spatial organization of pluripotency markers expression, we conducted mass spectrometry analysis using MALDI FTICR imaging, with a pixel size of 25 *µ*m and ultrahigh mass resolution to better track individual lipid species. We did not detect in these experiments changes in PC abundances. However, we observed a slight increase in *m/z* 940.5678 compared to the control, and an increase in *m/z* 742.5385 (PE 36:2) in days 5, 6, and 7, correlating with the changes in Oct4 expression in control versus the DZA-exposed sample. The most dramatic ion abundance increases compared to the control were for *m/z* 835.5346 (PI 34:1), 861.5499 (PI 36:2) and 863.5649 (PI 36:1) (Fig. 3).

**Table 1.** Sample sizes for analysis of temporal (A) and spatial (B) changes in phospholipid abundance.

| day | number of cells detected |  |  | number of dividing cells detected |  | number of edge cells detected |  |  |
| --- | --- | --- | --- | --- | --- | --- | --- | --- |
| | control | 35 μM | 100 μM | control | 35 μM | control | 35 μM | 100 μM |
| 0 | 8679 | 9571 | 9536 | 58 | 263 | 1481 | 864 | 2197 |
| 1 | 10587 | 19473 | 9681 | 223 | 458 | 1558 | 341 | 131 |
| 2 | 8129 | 11618 | 3190 | 66 | 102 | 1487 | 278 | 660 |
| 3 | 12483 | 14123 | 13321 | 119 | 299 | 1150 | 119 | 636 |
| 4 | 9156 | 12942 | 8214 | 48 | 365 | 485 | 322 | 1895 |
| 5 | 12295 | 18602 | 16051 | 151 | 292 | 466 | 621 | 101 |
| 6 | 9762 | 11655 | 9654 | 59 | 125 | 155 | 701 | 763 |
| 7 | 8105 | 16239 | 13800 | 112 | 373 | 477 | 407 | 613 |

**B**
| Cell count | control training |  | control validation |  |
| --- | --- | --- | --- | --- |
|  | TRA-181+ | SSEA-1+ | TRA-181+ | SSEA-1+ |
| day 6 | 1181 | 1252 | 4066 | 7286 |

| Cell count | 35 μM | | | 100 μM | |
| --- | --- | --- | --- | --- | --- |
|  | TRA-181+ | SSEA-1+ | NCAM-1+ | SSEA-1+ | NCAM-1+ |
| 5 | 1122 | 607 | 999 | 5226 | 6355 |
| 6 | 1470 | 3329 | 917 | 395 | 999 |
| 7 | 109 | 2537 | 2327 | 2525 | 4497 |

**Table 2.** Lipid annotations for 16 selected features found in all 3 spontaneous differentiation experiments. The table shows MALDI FTICR *m/z* values for the species of interest, proposed annotation, main adduct type detected, experimental monoisotopic *m/z* value, elemental formula, mass error (ppm), annotation confidence level, MS/MS collision energies, and fragment ions used to determine lipid annotation. The confidence level for lipid annotation was assigned as (1) exact mass, isotopic pattern, and MS/MS spectrum of a chemical standard matched to the feature. (2) exact mass, isotopic pattern, retention time, and MS/MS spectrum matched to an in-house spectral database or literature spectra (3) putative ID assignment based only on elemental formula match. (4) unknown compound. Asterisks (*) designate compounds for which fatty acid annotation was obstructed by contamination with fragments from a co-selected precursor ion. No matches for the head group of *m/z* 940.5678 species were found in METASPACE and LIPID MAPS databases.

| Precursor $m/z$ (FTICR) | Proposed Annotation | Adduct Type | Monoisotopic Mass | Ion Mass | Elemental Formula | Mass Error (ppm) | Confidence Level | Fragment Ions Used to Determine Annotation ($m/z$) | Collision Energy (eV) |
| --- | --- | --- | --- | --- | --- | --- | --- | --- | --- |
| 701.5122 | PA (18:1)/(18:0) and PA (20:1)/(16:0) | [M-H]<sup>-</sup> | 702.52 | 701.5127 | C<sub>39</sub>H<sub>75</sub>O<sub>8</sub>P | -0.68 | 2 | 255, 281, 283, 309, 419, 437 | 15 |
| 722.5129 | PE (P-16:0)/(20:4) | [M-H]<sup>-</sup> | 723.5203 | 722.513 | C<sub>41</sub>H<sub>74</sub>NO<sub>7</sub>P | -0.16 | 2 | 196, 259, 303, 419, 436 | 15 |
| 742.5385 | PE (18:1)/(18:1) | [M-H]<sup>-</sup> | 743.5465 | 742.5392 | C<sub>41</sub>H<sub>78</sub>NO<sub>8</sub>P | -0.98 | 2 | 281 | 15 |
| 748.5281 | PE O-38:6* | [M-H]<sup>-</sup> | 749.5359 | 748.5287 | C<sub>43</sub>H<sub>76</sub>NO<sub>7</sub>P | -0.76 | 3 | N/A | 15 |
| 778.5754 | PE (22:4)/(P-18:0) | [M-H]<sup>-</sup> | 779.5829 | 778.5756 | C<sub>45</sub>H<sub>82</sub>NO<sub>7</sub>P | -0.28 | 2 | 155, 331 | 20 |
| 819.5177 | PG (20:3)/(20:4) | [M-H]<sup>-</sup> | 820.5254 | 819.5182 | C<sub>46</sub>H<sub>77</sub>O<sub>10</sub>P | -0.56 | 2 | 227, 303, 305 | 20 |
| 821.5331 | PG (18:1)/(22:5) | [M-H]<sup>-</sup> | 822.5411 | 821.5338 | C<sub>46</sub>H<sub>79</sub>O<sub>10</sub>P | -0.87 | 2 | 227, 281, 329 | 20 |
| 833.5181 | PI (18:1)/(16:1) | [M-H]<sup>-</sup> | 834.5258 | 833.5186 | C<sub>43</sub>H<sub>79</sub>O<sub>13</sub>P | -0.55 | 2 | 223, 241, 253, 281 | 25 |
| 835.5346 | PI (16:0)/(18:1) | [M-H]<sup>-</sup> | 836.5415 | 835.5342 | C<sub>43</sub>H<sub>81</sub>O<sub>13</sub>P | 0.47 | 2 | 223, 241, 255, 281 | 25 |
| 859.5342 | PI (16:0)/(20:3) | [M-H]<sup>-</sup> | 860.5415 | 859.5342 | C<sub>45</sub>H<sub>81</sub>O<sub>13</sub>P | 0 | 2 | 223, 241, 255, 305 | 20 |
| 861.5499 | PI (18:1)/(18:1) | [M-H]<sup>-</sup> | 862.5571 | 861.5499 | C<sub>45</sub>H<sub>83</sub>O<sub>13</sub>P | 0.05 | 2 | 223, 241, 281 | 25 |
| 863.5649 | PI (18:1)/(18:0) | [M-H]<sup>-</sup> | 864.5728 | 863.5655 | C<sub>45</sub>H<sub>85</sub>O<sub>13</sub>P | -0.7 | 2 | 223, 241, 281, 283 | 25 |
| 883.5336 | PI (18:1)/(20:4) | [M-H]<sup>-</sup> | 884.5415 | 883.5342 | C<sub>47</sub>H<sub>81</sub>O<sub>13</sub>P | -0.68 | 2 | 223, 241, 281, 303 | 25 |
| 885.5494 | PI (18:0/20:4) | [M-H]<sup>-</sup> | 886.5571 | 885.5499 | C<sub>47</sub>H<sub>83</sub>O<sub>13</sub>P | -0.51 | 2 | 223, 241, 283, 303 | 25 |
| 911.5661 | PI 40:5* | [M-H]<sup>-</sup> | 912.5728 | 911.5655 | C<sub>49</sub>H<sub>85</sub>O<sub>13</sub>P | 0.65 | 3 | N/A | 20 |
| 940.5678 | Unknown | [M-H]<sup>-</sup> | N/A | N/A | N/A | N/A | 4 | N/A | 35 |

**Figure 3.**
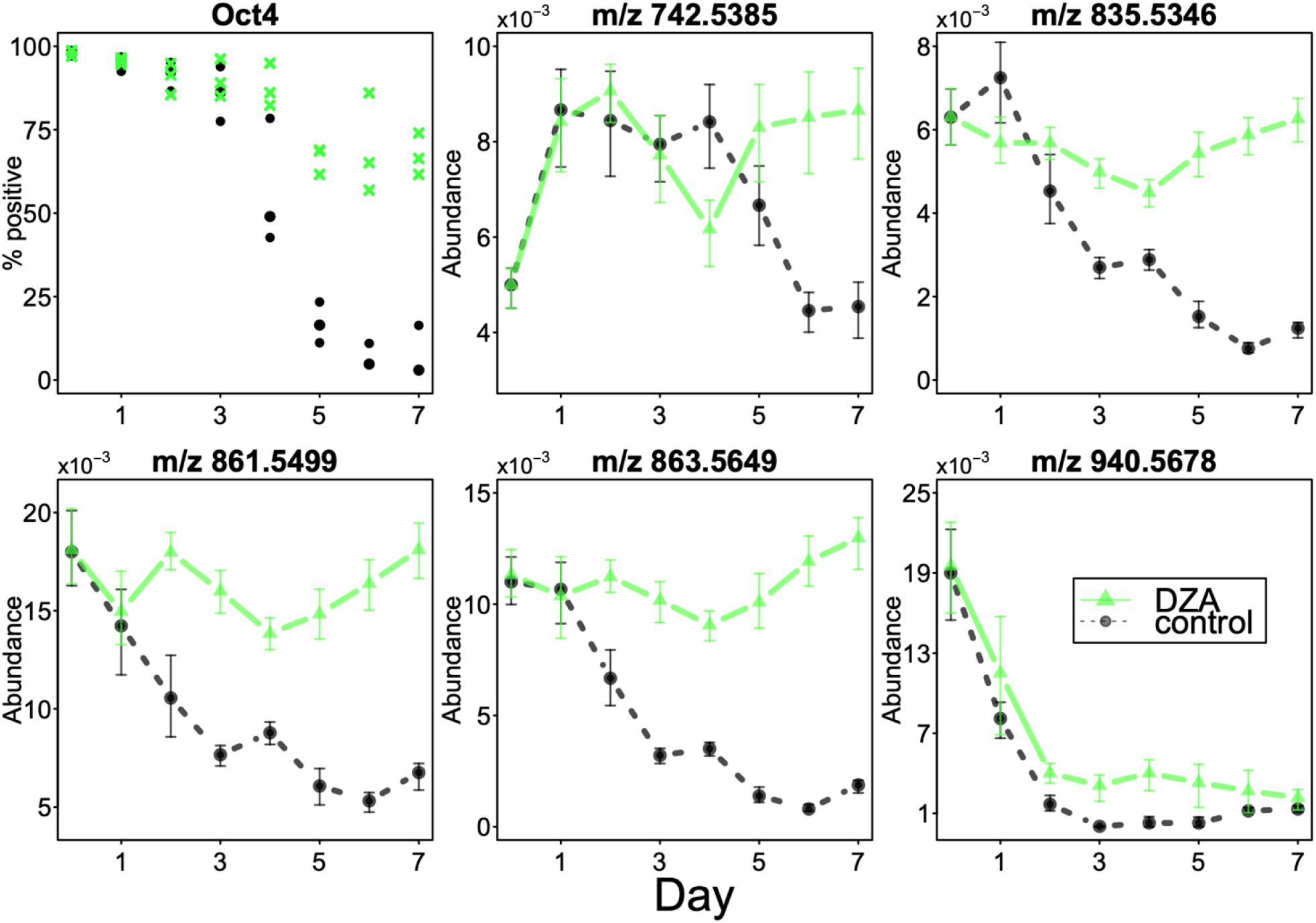
Continuous exposure to 3-Deazaadenosine (DZA) promotes pluripotency maintenance through perturbation of phospholipid levels. More than 50% of population maintained Oct4 expression in the DZA-exposed sample in 3 independent experiments (top left). MALDI FTICR MS analysis of control and DZA-exposed samples revealed that several phospholipids that decline with differentiation time in the control experiment maintain their levels in the DZA experiment, correlating with the Oct4 expression. Data points represent average *m/z* abundances per image, error bars show 25^th^ and 75^th^ percentiles.

### Inhibition of phosphatidylinositol 3-kinase results in increased NCAM-1 expression and changes in colony spatial organization

The *m/z* 835.5346, 861.5499 and 863.5649 species detected by MALDI FTICR MS belong to the phosphatidylinositol (PI) family (Table 2). To further clarify the importance of PI cycling in the differentiation process, we conducted a series of experiments in which we initiated differentiation while inhibiting phosphatidylinositol 3-kinase with LY294002. We grew 8 iPSC colony samples, one for each day of spontaneous differentiation, with a low inhibitor concentration of 35 *µ*M (Fig. 4a) and another 8 samples with a high inhibitor concentration of 100 *µ*M (Fig. 4b). While performing confocal microscopy imaging on these samples, we observed a dose-dependent increase in NCAM-1 expression compared to controls, as well as changes in spatial organization of NCAM-1 and SSEA-1 positive cells (Fig. 4a, b, Fig. 5). When comparing phospholipid abundance trajectories between the 3 conditions (control, 35 *µ*M and 100 *µ*M inhibition, Fig. 4c), we observed the absolute value of trajectories slopes increase in a dose-dependent manner for several PI family members (*m/z* 859.5, 863.5, 883.6 and 911.5). The representative *m/z* 748.5 ion showed consistent growth in all three conditions as well as a spatial correlation with SSEA-1 expression and a strong anti-correlation with NCAM-1 expression (Fig. 4a, b). The distinctive trajectories of *m/z* 778.5 and 940.6 were conserved with PI 3-kinase inhibition. We observed that cells remained more pluripotent on the edge of the colony over the course of differentiation from immunocytochemistry performed on iPSC colonies stained with Oct4 for pluripotency, Otx2 for ectoderm differentiation, and Pax6 for neural lineage (Fig. 6 supp. 1a). To compare phospholipid abundances in the center and on the edge of the colony, we divided cells into 7 groups based on their location in the colony and calculated average *m/z* ion abundances for 7 different distances from the edge. Some phospholipids (*e*.*g. m/z* 722.5 and 748.5) gradually increased in abundance with distance from the edge and some gradually decreased (*e*.*g. m/z* 778.5 and 940.6), mostly consistent with their previously shown pluripotency correlation. Examples of such trends for day 3 in the control experiment are shown in Fig. 6a. Immunocytochemistry images (Fig. 6 supp. 1a) suggested that the difference between the edge and the center of the colony became more prominent with overall colony differentiation, consistent with some phospholipids showing a higher correlation with edge distance in later days of differentiation and little correlation on day 0 (Fig. 6b). We also observed a correlation “flip” for some lipids (*e*.*g*., *m/z* 940.6) in the PI 3-kinase inhibited experiment (Fig. 6b). While this trend is not reflected in immunocytochemistry images of days 0-3 of the PI 3-kinase inhibited differentiation, day 4 starts to reveal a mixed Oct4/Otx2 pattern, with days 5 and 6 in the 100 *µ*M LY294002 experiment showing a reversed spatial pattern of pluripotency, with increased Otx2 expression on the edge of the colony and Oct4 expression in the center (Fig. 6 supp. 1b).

**Figure 4.**
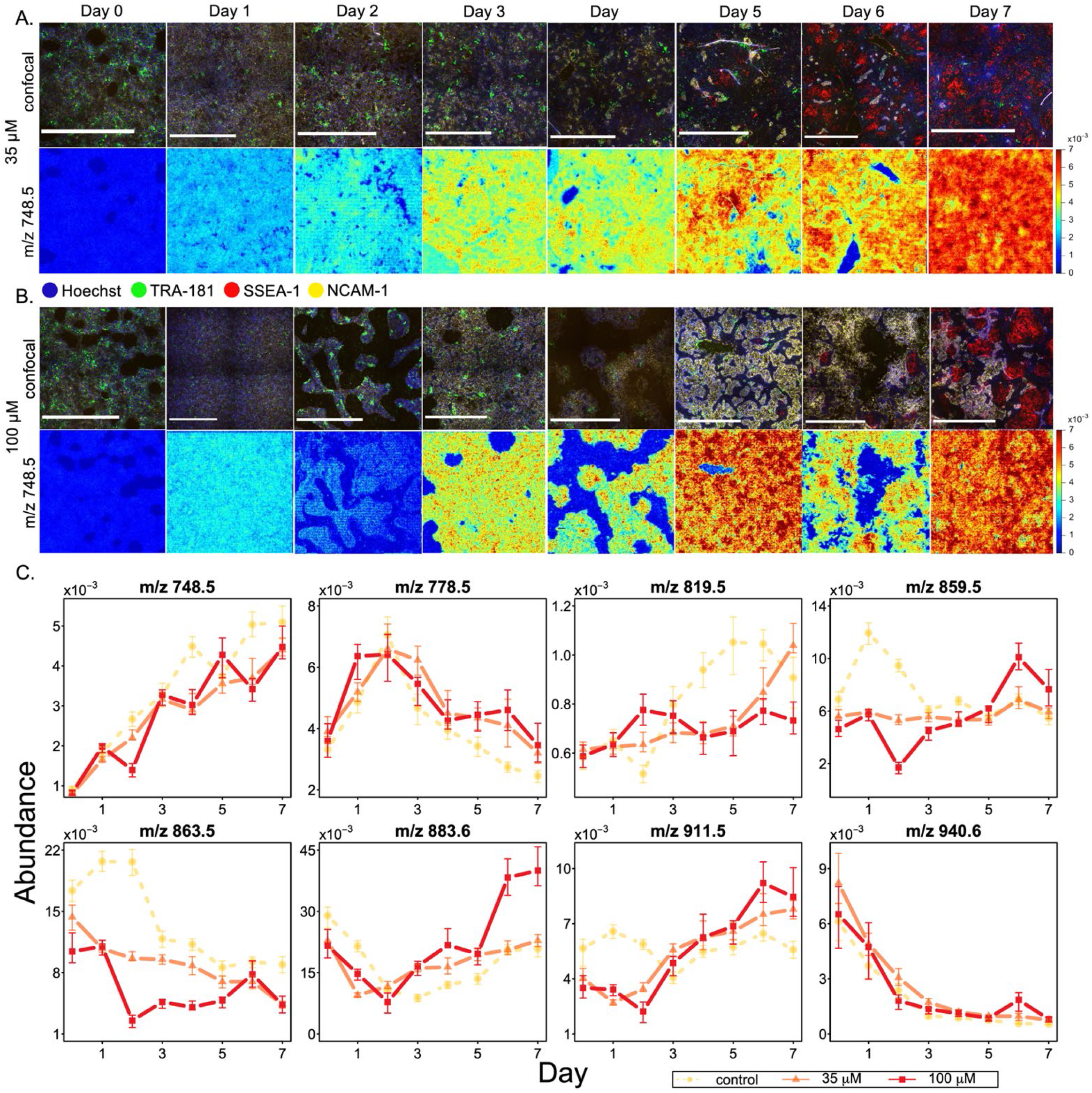
Spontaneous differentiation with phosphatidylinositol 3-kinase inhibition as observed by phospholipid levels via MALDI imaging. A, B. Top row – confocal images of iPSC colonies undergoing differentiation for 7 days with addition of LY294002 on day 0, blue is Hoechst staining, green is TRA-181, red is SSEA-1, yellow is NCAM-1. Bottom row shows corresponding MALDI ion images for *m/z* 748.5 with blue colors representing low peak abundance and red representing high abundance. C. Temporal changes in mean phospholipid abundance during spontaneous differentiation based on the LY294002 dose. Error bars show 25^th^ and 75^th^ percentiles.

**Figure 5.**
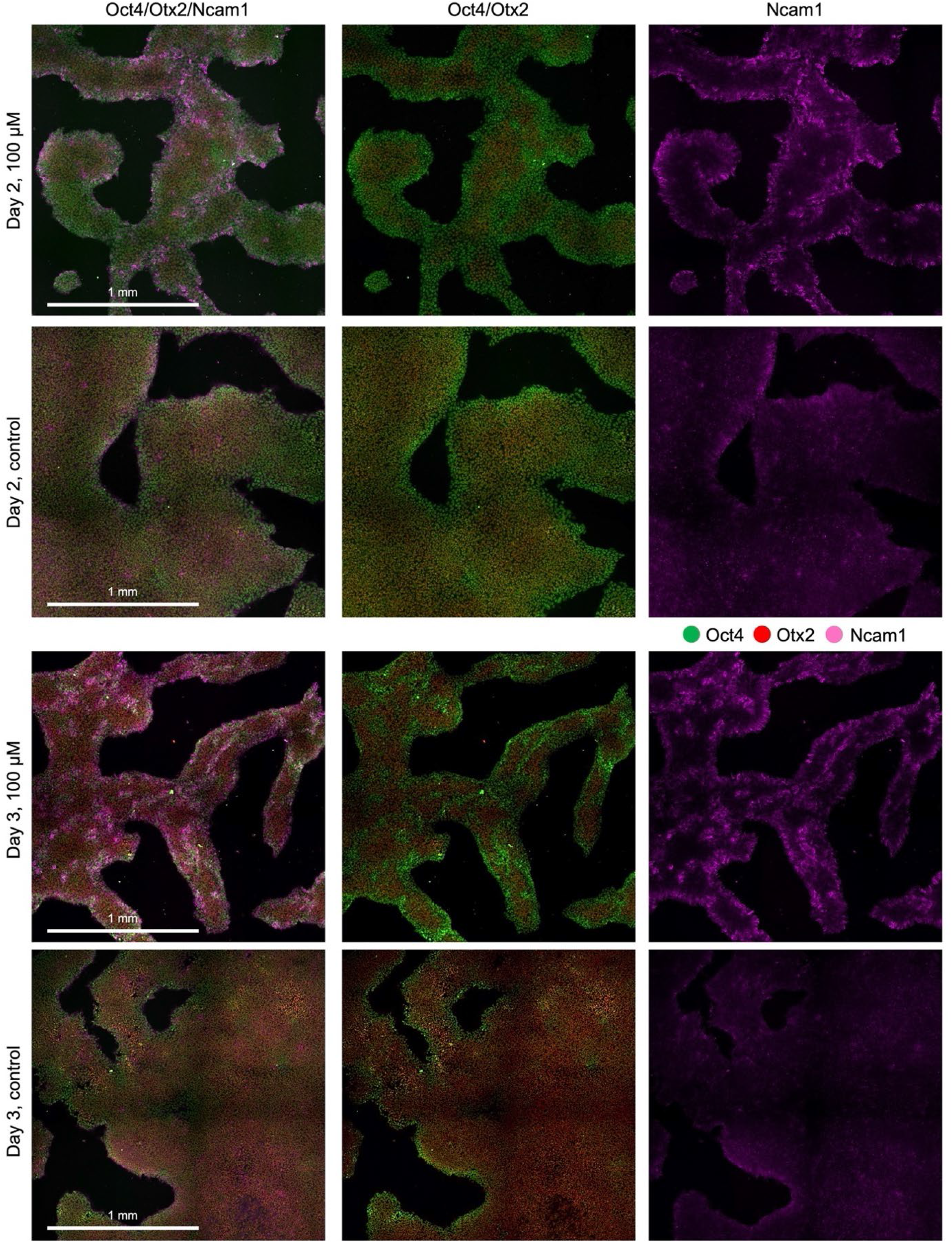
Ncam1 and Oct4 spatial expression in iPSC colonies are altered with LY294002. Edge-independent patterns of Ncam1/Oct4 expression from immunoflourescence imaging are observed as early as day 2 and 3 of PI 3-kinase inhibited differentiation with 35 µM or 100 µM LY294002, in contrast to the control differentiation with vehicle. Scalebars are 1 mm.

**Figure 6.**
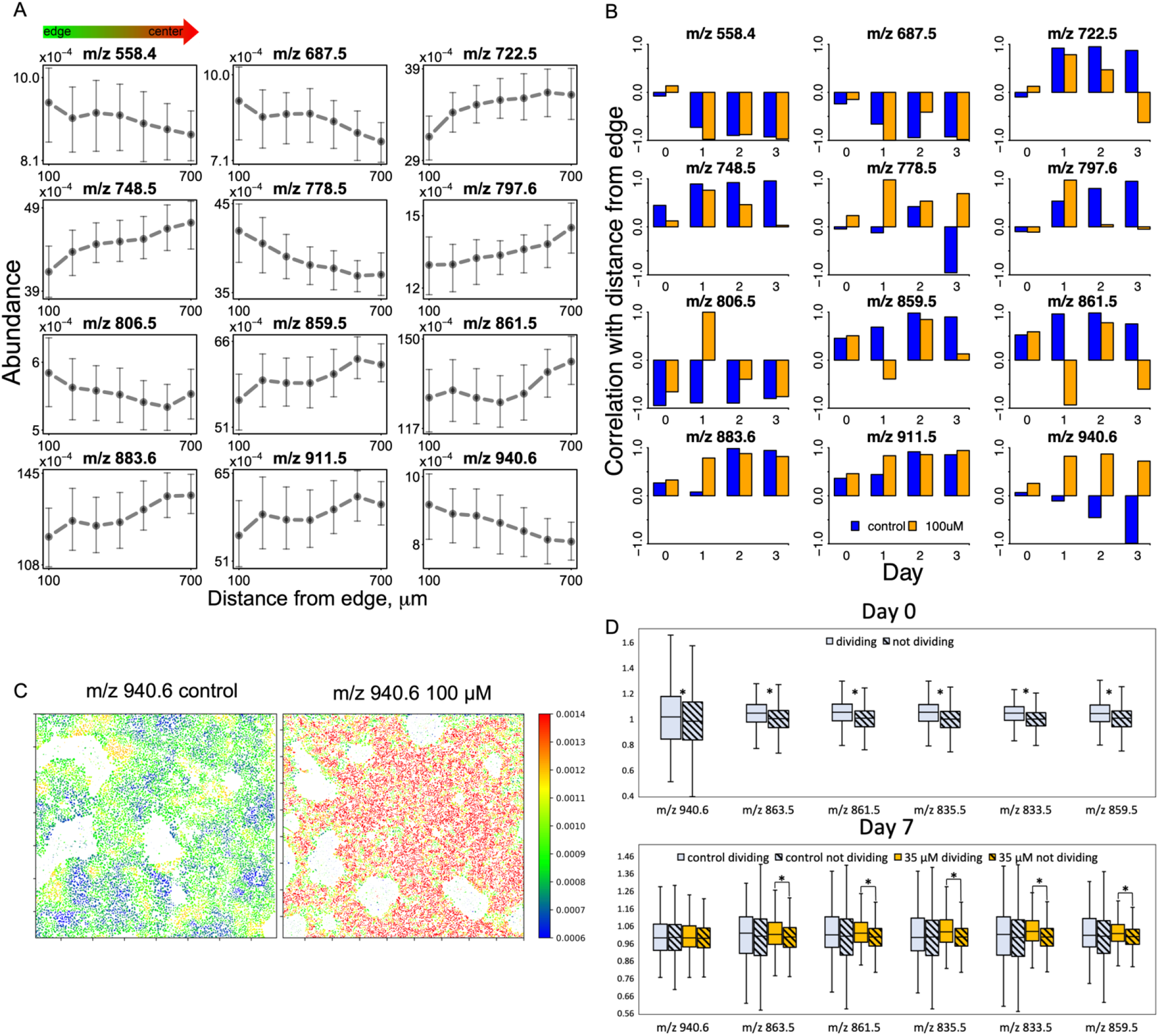
Phospholipid abundances change with colony and cell morphology. A. Mean phospholipid abundances in day 3 of control spontaneous differentiation change with the distance from the edge of the colony. Points represent mean values within the 100 μm edge distance range, error bars show 25 and 75 percentiles. B. Correlation of phospholipid abundances with edge distance changes with days of differentiation and with LY294002 addition. For illustration of edge dependent expression of Oct4 and Otx2 refer to Figure 6 – figure supplement 1. C. Spatial distribution of *m/z* 940.6 abundance in day 3 of the control experiment shows increased signal on the edge of the colony in contrast with high LY294002 dose experiment, which shows decreased signal on the edge of the colony. D. Differences in neighbor-relative lipid abundances in dividing versus non-dividing cells. For of manually annotated validation dataset refer to Figure 6 – figure supplement 2. Top: presented lipids are significantly more abundant in dividing cells in day 0. Bottom: by day 7 control samples stop exhibiting significant differences in lipid abundances, while differences in PI 3-kinase inhibited samples are still significant. Shaded boxes represent non dividing cells. Box boundaries show 25^th^ and 75^th^ percentiles, middle line shows median, whiskers show minimum and maximum values. Asterisks shows statistical significance in median differences (p-value < 0.05), for exact p-values refer to Figure 6 – Source data 1.

### Phospholipid abundances vary based on cell’s proliferative status

Because PI 3-kinase signaling is strongly related to cell proliferation, we hypothesized that cells undergoing mitosis would reflect differences in PI signatures. To find metabolic signatures corresponding to mitotic cells, we developed a k-means clustering algorithm to distinguish cells undergoing mitosis by their nuclear morphology and the brightness of Hoechst stain. To test the algorithm, we manually annotated dividing nuclei in a small ROI; the algorithm yielded 98.8% prediction accuracy. Next, using overlaid and aligned MALDI MS images, we associated the cell’s proliferative status to its metabolic signature. Since this task required precise single-cell comparison, we used a neighbor-relative abundance metric to account for potential unevenness of the background. Finally, we compared the ion abundances between dividing and non-dividing cells on day 0 of differentiation (Fig. 6d, top). We observed higher neighbor-relative abundances from *m/z* 835.5, 861.5, 863.5 and 940.6 in dividing cells, which is consistent with these ions’ previous correlations with pluripotency. By day 7 of the control differentiation these differences disappear; however, they are maintained in the PI 3-kinase inhibited differentiation (Fig. 6d, bottom). Statistical significance was determined by the two-tailed Mann-Whitney U test with significance threshold of p-value < 0.05. All resulting statistics are summarized in Figure 6 – Source data 1.

### Phospholipid profiles reveal a bifurcation in cell lineage specification upon PI 3-kinase inhibition

The mixed phenotypes that emerged from LY294002 treatment (Fig. 7a) suggested a potential progression between multiple cell populations that could be spatiotemporally investigated further. To quantify spatial correlation of lipid abundances and fluorescent labels in PI 3-kinase inhibited experiments and identify metabolic signatures corresponding to newly emerging cell populations, we trained two PLS-DA classifiers, one for each dose of inhibitor, using the last three days of differentiation. With 35 *µ*M of LY294002 we observed 3 distinct cell populations: SSEA-1 positive (SSEA-1+), NCAM-1 positive/TRA-181 positive (NCAM-1+), and NCAM-1 negative/TRA-181 positive (TRA-181+). Since day 6 prominently exhibits all 3 populations, we trained a model on the day 6 image, and withheld data from day 5 and day 7 to use as validation sets. After variable trimming training set yielded 82% accuracy, day 5 and day 7 yielded 66% and 71% accuracy respectively (Fig. 7 – supp. 1a). The PLS-DA biplot (Fig. 7 – supp. 1b) shows distinct SSEA-1+ and NCAM-1+ correlated clusters of both observations (scores) and variables (loadings). These clusters of variables represent distinct lipid signatures of the two populations: the SSEA-1+ population had increased levels of *m/z* 722.5, 748.5, 819.5, and *m/z* 821.5 while the NCAM-1+ population had increased levels of PI lipids (*m/z* 859.5, 861.5, 863.5, 883.5, 885.6), along with *m/z* 778.5 and 940.6. We also observed that the TRA-181+ cluster is an intermediate between two other clusters, possibly indicating these cells as not yet committed to a particular lineage. When this analysis was repeated with only SSEA-1+ and NCAM-1+ cells, the training accuracy improved to 95%; day 5 and day 7 yielded 85% and 97% accuracy respectively. With 100 *µ*M of LY294002 we only observed 2 distinct cell populations: SSEA-1+ and NCAM-1+, and no TRA-181 positive cells, suggesting that higher concentration of inhibitor pushes cells towards commitment to a particular lineage. Since day 7 has equal representation of both populations, we used it as a training set and withheld day 5 and day 6 as validation sets. After variable trimming the training set yielded 95% accuracy, day 5 and day 6 yielded 80% and 90% accuracy respectively (Fig. 7b). Finally, PLS-DA biplot for the 100 *µ*M condition (Fig. 7c) revealed similar metabolic signatures of the two populations as in the 35 *µ*M condition, showing that the selected lipids and pluripotency markers correlate consistently between days and inhibitor concentrations.

**Figure 7.**
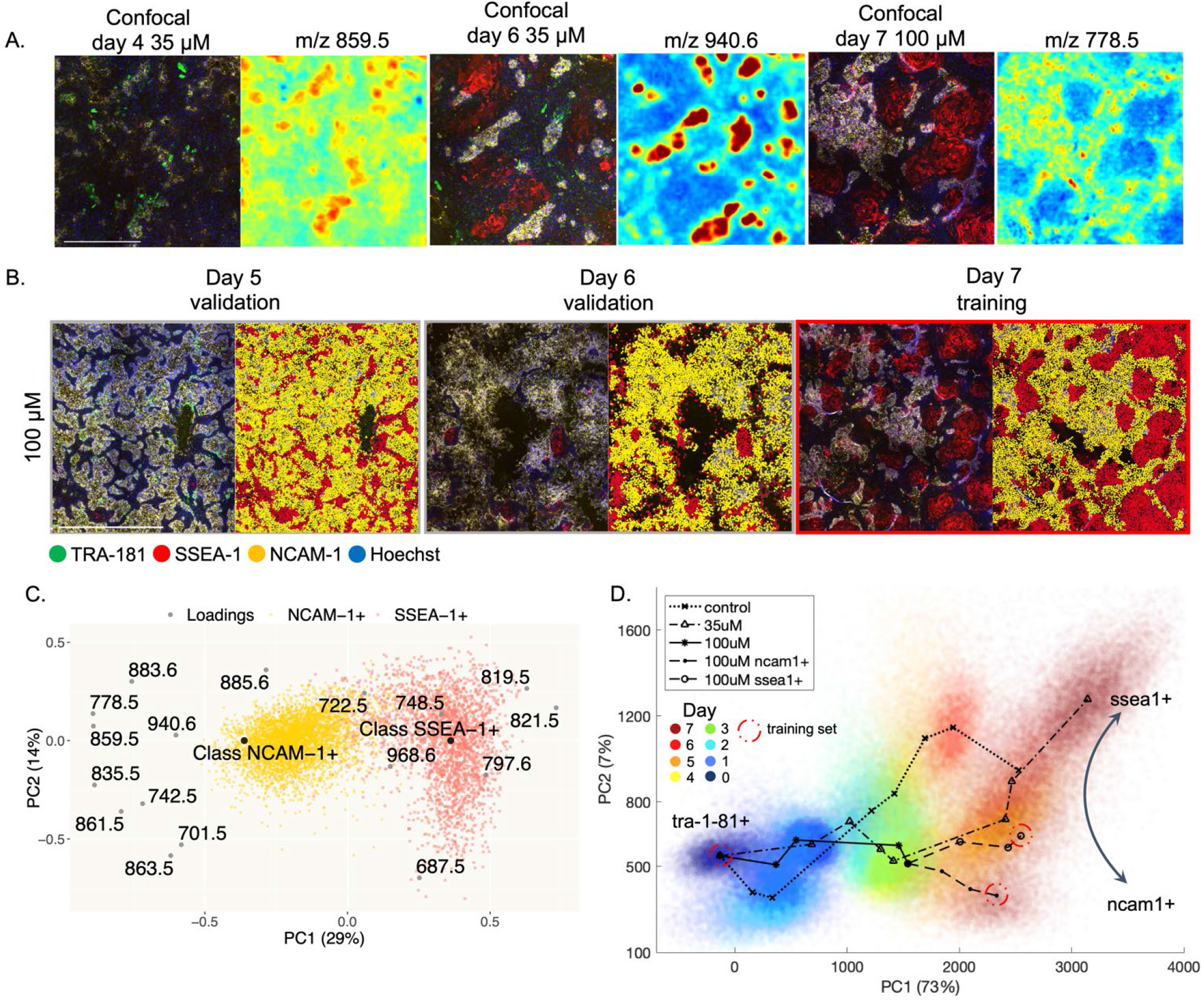
Changes in spatial organization of pluripotency markers and phospholipids with PI 3-kinase inhibition are predictable by multivariate analysis. A. Examples of ionic species correlating with cell lineage markers. Colors in confocal images are as follows: blue is Hoechst, green is TRA-181, red is SSEA-1, yellow is NCAM-1. MALDI ion images are pseudo colored with blue showing low abundances and red showing high abundances. Scalebar is 0.5 mm. B. Confocal images of days 5-7 of spontaneous differentiation with 100 µM of LY294002 and their predicted lineage labels. Red color labels SSEA-1+ cells, yellow colors NCAM-1+ cells. Day 7 was used as a training set, days 5 and 6 as validation sets (80% and 90% accuracies). Scalebar is 1 mm. The data for the 35 µM experiment can be found in Figure 7 – figure supplement 1. C. Biplot of the PLS-DA model used to discriminate between the cell populations in Fig. 7b. D. A principal component space created by training a PLS-DA model with 3 main populations: pluripotent cells (day 0), NCAM-1+ and SSEA-1+ cells of day 7 (100 µM of LY294002). The rest of the data from all 3 experiments was projected into this principal component space. Red colors indicate later days of the differentiation, blue colors indicate early days.

To summarize the relationships between all 3 experiments, we selected 3 main observed phenotypes as a training set for a PLS-DA model: the SSEA-1+ and NCAM-1+ populations from day 7 of the 100 *µ*M condition and cells from day 0 as a pluripotent population. Next, we projected all the data into that principal component space (Fig. 7d). We observed a correlation of day of differentiation and PC1, indicating that PC1 represents time in principal component space. We also observed the divergence of NCAM-1+ and SSEA-1+ populations for days 5-7 of the 100 *µ*M condition, with the 35 *µ*M and control trajectories corresponding to the SSEA-1+ branch, suggesting that PC2 is representative of cell fate. A compilation of our findings is provided for phospholipid species consistently connected to cell fate throughout our analysis (Fig. 7 – supp 1c).

## Discussion

Induced pluripotent stem cells are emerging as a powerful regenerative medicine tool for the creation of patient-specific tissues for autologous transplantation [15]. Investigating mechanisms underlying an initial loss of pluripotency in iPSCs is desirable for revealing early quality control targets, preventing the wasting of time and resources on a batch bound to fail ([16], [17]). We exploited and enhanced a previously developed multimodal image analysis approach [18] to establish the temporal sequence of metabolic changes associated with differentiation and relate these changes to pluripotency surface markers and the spatial distribution of phospholipid abundances within the cell colony. This approach allowed us to establish robust and predictable early metabolic markers of pluripotency loss during spontaneous differentiation that provide insight into the heterogeneity of iPSC populations; because these changes occur earlier than decline in Oct4 expression, these phospholipids hold potential as quality control targets in cell manufacturing. Our co-registration image analysis on iPSC spontaneous differentiation revealed that *m/z* 742.5 (PE 36:2), 778.5 (PE 22:4), 835.5 (PI 34:1), 859.5 (PI 36:3), 861.5 (PI 36:2), 863.5 (PI 36:1) and 940.6 spatially correlated to the TRA-181 surface pluripotency marker and decreased with differentiation time, consistent with the decrease in Oct4 expression. This provided evidence that these phospholipids could be used as early metabolic markers of differentiation.

Informed by MALDI MS experiments and literature searches, we inhibited PEMT activity during spontaneous differentiation by continuous cell exposure to DZA, which resulted in sustained maintenance of pluripotency as confirmed by elevated Oct4 expression. We consistently observed PI lipids (*m/z* 835.5, 861.5, 863.5) preceding changes in Oct4 expression which, once again, suggests that metabolic changes may occur earlier than changes in transcription factors. MicroRNAs have been reported as master metabolic controllers of naïve to primed ESC state and reprogramming to iPSCs, and potentially are altering lipid enzyme expression levels in advance of differentiation in iPSCs ([19], [20]). While the unknown species at *m/z* 940.6 did not match changes in Oct4 expression that occur with DZA inhibition, it could reflect an underlying spectrum of cell pluripotency status – including epigenetic changes - that precede the drop in Oct4 expression. This lipid species resisted all attempts of structural annotation due to its comparatively lower signal-to-noise ratio, even with some of the most modern MALDI imaging MS instrumentation available and extensive MS/MS analysis attempts.

Since several of these discovered lipid markers belong to the PI family, we hypothesized that PI cycling regulates the differentiation process. Supporting this hypothesis, we report that PI 3-kinase inhibition during spontaneous differentiation results in increased neural lineage specification and distinct spatial clustering of cells with similar cell fate marker expression. Such clustering further highlighted the spatial correlation of certain phospholipids and cell lineage markers. The ion at *m/z* 748.5 (PE O-38:6) was strongly anti-correlated with NCAM-1 expression and correlated with SSEA-1 expression, as well as consistently increasing with differentiation time in all 3 experiments. The unknown lipid species at *m/z* 940.6 strongly correlated with NCAM-1 expression, along with other metabolic markers that correlated with pluripotency in previous experiments. These findings suggest that observed NCAM-1 positive cell population is metabolically closer to the pluripotent state than the rest of the colony, which we confirmed by immunocytochemistry images showing co-expression of NCAM-1 and Oct4 in PI 3-kinase inhibited experiments (Fig. 5).

Our metabolic imaging enabled new insights in the role subpopulations of cells are playing within a cell colony. Cells remained more pluripotent on the edge of the colony compared to the colony center throughout the spontaneous differentiation; however, in certain cases this pattern was disrupted with PI 3-kinase inhibitor treatment due to spatial reorganization of the distinct cell lineages. MS and confocal imaging co-registration allowed us to compare metabolic states of cells on the edge of the colony with cells in the colony center. Under control conditions, we consistently observed phospholipids correlating with differentiation to be more abundant in the colony center, and species corresponding to pluripotent cells to have higher levels on the edge of the colony. However, in the experiments treated with a high dose of PI 3-kinase inhibitor, we observed these correlations to be disrupted, and for some species (e.g., *m/z* 940.6) we observed a reversed trend. PI 3-kinase activates Akt which is involved in cell migration and mTOR pathways, perhaps explaining formation of spatial clusters of lineage markers in an edge-independent way in PI 3-kinase inhibited colonies – possibly, the spontaneous centers of differentiation do not migrate out into the colony, creating a more localized progeny.

Finally, the high resolution of confocal images allowed not only segmentation of individual nuclei but also identification of their mitosis stage; in combination with MALDI MS images, it facilitated a connection with each cell’s metabolic profile. This analysis highlighted phospholipids shown to correlate with pluripotency to be significantly more abundant in dividing cells on day 0, consistent with the faster cell cycle of pluripotent cells. Interestingly, by day 7 this difference disappears in the control experiment; however, it is maintained in PI 3-kinase inhibited samples. Akt is also involved in cell proliferation - possibly, cells that continue to divide despite PI 3-kinase inhibition have a more contrasting phenotype compared to dividing cells in the control condition.

Our multi-modal imaging co-registration pipeline produced robust datasets that tied together cells’ location, morphology, cell fate surface markers and metabolic profile. Multivariate analysis performed on these datasets consistently illustrated the predictive power of metabolic data, allowing for accurate prediction of priming for differentiation or a cell’s surface marker expression. Our analysis also informed multivariate trajectories revealing divergent metabolic cell fate, which could be useful in a manufacturing setting by identifying key windows of differentiation in which lineage specification can be manipulated and/or corrected. Future work includes further investigation of the role of phosphatidylinositols in self-organization of 3D iPSC organoids and elucidation of additional label-free morphological features that reflect the lipid signatures discovered here for cell manufacturing applications.

## Acknowledgements

This material is based upon work supported by the National Science Foundation under Grant No. EEC-1648035. We gratefully acknowledge support from the NSF CMaT NSF Research Center and the Marcus Center for Therapeutic Cell Characterization and Manufacturing. The authors also acknowledge support from NSF MRI CHE-1726528 grant for the acquisition of an ultra-high resolution Fourier transform ion cyclotron resonance (FTICR) mass spectrometer for the Georgia Institute of Technology core facilities. The authors also thank Dr. Li Li for initial MS experiments in early stages of this project.

## Author contributions

MLK, FMF, AAN and AVG conceived the idea and designed the studies. AVG, AAN and DH carried out the experiments. AAN and TR performed the computational analysis of the results. AAN, MLK and FMF wrote the manuscript.

## Data Availability

The datasets generated and/or analyzed during the current study are available from the corresponding author upon request.

## Code Availability

The co-registration and segmentation code with data samples can be found at our lab’s GitHub (https://github.com/kemplab/coregistration_gui).

## Methods

### S1. Materials

Acetonitrile (LC-MS grade), ammonium formate, and norharmane (98%) were purchased from Fisher Chemical (Pittsburgh, PA, USA). Laboratory grade Triton™ X-100, 3-Deazaadenosine, acetone and red phosphorus (≥99.99% purity) were purchased from Sigma Aldrich (Sigma-Aldrich Corporation, St. Louis, MO, USA). Ultrapure water with 18.2 MΩ·cm resistivity (Barnstead Nanopure UV ultrapure water system, USA) was used to prepare the ammonium formate buffer wash solution. Conductive ITO glass slides were purchased from Bruker Daltonics (Billerica, MA, USA). Dow SYLGARD™ 184 Silicone Encapsulant Clear Kit was purchased from Ellsworth Adhesives (Loganville, GA, USA). Anhydrous DMSO, Corning® Matrigel® Growth Factor Reduced (GFR) Basement Membrane Matrix (Phenol Red-Free, *LDEV-Free), KnockOut™ DMEM, Dulbecco’s Phosphate-Buffered Saline, 10X with calcium and magnesium (DPBS), B-27™ Supplement (50X), Molecular Probes Hoechst 33342, and Donkey anti-Mouse IgG (H+L) Highly Cross-Adsorbed Alexa Fluor Plus 488 Secondary Antibody were purchased from Thermo Fisher Scientific (Waltham, MA, USA). MTeSR™ Plus (Basal Medium and 5X Supplement), Y-27632 RHO/ROCK pathway inhibitor, and accutase were purchased from Stemcell Technologies (Cambridge, MA, USA). RPMI 1640 was purchased from Caisson Labs (Smithfield, UT, USA). GloLIVE Human Pluripotent Stem Cell Live Cell Imaging Kit (catalog #NLLC2155R) including positive marker TRA-1-81 and negative marker SSEA-1 (301021), and Alexa Fluor 647-conjugated Human NCAM-1/CD56 (MC-480) live stain (catalog #FAB24081R-100UG) were purchased from R&D Systems (Minneapolis, MN, USA) as well as Human Three Germ Layer 3-Color Immunocytochemistry Kit (catalog #SC022) including Anti-Human Otx2 NL557-Conjugated Goat IgG. 6-Well Tissue Culture Plates were purchased from Celltreat (Pepperell, MA, USA). Paraformaldehyde Aqueous Solution was purchased from Electron Microscopy Sciences (Hatfield, PA, USA). Odyssey® PBS Blocking Buffer was purchased from LI-COR Biosciences (Linkoln, NE, USA). Mouse Anti-human Oct-3/4 (C-10) primary antibody (catalog #sc-5279) was purchased from Santa Cruz Biotechnology (Dallas, TX, USA). LY294002 was purchased from Cell Signaling Technology (Danvers, MA, USA). Mouse anti-Human Alexa Fluor 647-conjugated Pax6 antibody (O18 1330) was purchased from BD Biosciences (San Jose, CA, USA, catalog #562249).

### S2. Cell culture

HiPSC WTC11 cells (Coriell Institute, catalog ID GM25256, sex: male) were grown in 6-well plates. Wells were coated with 1mL Matrigel (GFR in Knockout D-MEM, 1:100) per well and incubated overnight. Cells were fed 2 mL media per well daily (basal MTeSR Plus media + supplement, 4:1). During passage, 0.5 mL accutase was added to each well and incubated for 3 min. Cells were lifted and collected into a 15 mL tube with excess DPBS (∼3x times accutase used). Cells were centrifuged at 1000 rpm for 5 minutes, supernatant was removed, and the pellet was lifted in 2 mL media with 2 µL Rock inhibitor. Cells were seeded at 100-200 K density in 2 mL media and 2 µL Rock inhibitor was added for the first day after passage, then cells were fed as usual. To start spontaneous differentiation media was switched to RPMI plus B-27 Supplement (49:1) the next day after seeding. In experiments with PI 3-kinase inhibition LY294002 powder was reconstituted at 25 mM in DMSO and added to the RPMI/B-27 media at 35 or 100 µM during the first 24 hours of spontaneous differentiation, after which cells were fed fresh RPMI/B-27 media. Equal amount of pure DMSO was added to a corresponding control experiment. In experiments with PEMT inhibition 3-Deazaadenozine powder was reconstituted at 50 mM in DMSO and added daily to fresh RPMI/B-27 media during feeds at 50 µM. Equal amount of pure DMSO was added daily to a corresponding control experiment.

### S3. Flow cytometry

Eight control wells and 8 DZA-exposed wells underwent 0 to 7 days of spontaneous differentiation in 6-well plates, each well was reproduced 3 times. Cells were lifted with 0.5 mL accutase per a well of a 6-well plate, centrifuged at 1000 rpm for 5 minutes, supernatant was removed, and the pellet was lifted in 1 mL of 4% paraformaldehyde solution in PBS for cell fixation. After 10 min at room temperature cells were centrifuged again and resuspended in 1 mL of 0.3% Triton™ X-100 solution in PBS for permeabilization. After 15 min at room temperature cells were centrifuged and resuspended in 1 mL Odyssey Blocking Buffer for 1 hour at room temperature. Next, cells were centrifuged and resuspended in 1 mL Odyssey Blocking Buffer with 5 µL mouse anti-human Oct4 primary antibody and were left at 4C overnight. After that, cells were centrifuged and resuspended in 1 mL PBS as a wash step. At this point, 0.5 µL of cells from Day 0 sample were set aside as a negative control. Next, cells were centrifuged and resuspended in 1 mL Odyssey Blocking Buffer with 1 µL anti-mouse Alexa Fluor Plus 488 secondary antibody for 30 minutes in the dark. Finally, after another wash step, cells were centrifuged and resuspended in 1 mL PBS and transferred to a FACS tube through the strainer cap. Samples were analyzed on BD FACSMelody™ Cell Sorter (BD Biosciences, San Jose, CA, USA) with excitation wavelength of 488 nm, detection in 515 nm-545 nm range. Data acquisition was performed with BD FACSChorus™ Software (BD Biosciences), data processing was performed with FlowJo™ (BD Biosciences). Oct4 gate was created so 99.9% of negative control fall into Oct4-negative category. Percentage of cells registered as Oct4-positive according to the gating was recorded for each sample.

### S4. Immunocytochemistry

Eight control wells and 8 wells exposed to 100 µM LY294002 underwent 0 to 7 days of spontaneous differentiation on ITO-covered glass slides with a glued PDMS 8-well wall. Wells were washed with 100 µL of PBS each (wash step) and then fixed with 100 µL of 4% paraformaldehyde solution in PBS for 10 minutes. After 3 wash steps cells were permeabilized for 15 minutes with 100 µL of 0.3% Triton™ X-100 solution in PBS. After a wash step, cells were blocked with 100 µL of Odyssey Blocking Buffer for 1 hour at room temperature. Next, we diluted mouse anti-human Oct4 primary antibody at 1:200 ratio in Odyssey Blocking Buffer, added anti-human NL557-Conjugated Otx2 antibody at 1:100 ratio as well as anti-human Alexa Fluor 647-conjugated Pax6 antibody at 1:50 ratio. Cells were treated with 100 µL of antibody mixture for 1 hour in the dark. Next, after 3 wash steps cells were treated with 100 µL of Odyssey Blocking Buffer with Hoechst (1:1000) and anti-mouse Alexa Fluor Plus 488 secondary antibody (1:1000) for 30 minutes in the dark. Finally, after 3 wash steps 100 µL of PBS was added to each well and cells were imaged with Nikon UltraVIEW VoX W1 Spinning Disk Confocal with sCMOS camera at 10x magnification (0.65 mm/px), 100 ms exposure and 100% laser power for all wavelengths.

### S5. Co-registration Sample Preparation

SYLGARD™ was poured in a custom-made 3D-printed molds and placed in the 70°C oven for 3 hours. The resulting 8-well silicone wall was adhered to an Indium-Tin-Oxide (ITO)-coated slide with SYLGARD™ and was placed in the oven for 30 min. Next, for 8 consecutive days cells were seeded in the corresponding wells: 1 well per day was coated with 100 µL of Matrigel and incubated for 1 hour, then hiPSC WTC11 cells were seeded onto it at 2000 cells/*mm*^2^ density. On the second day after seeding in each well the spontaneous differentiation was initiated. By day 9 of repeating these steps the slide contained samples of 8 consecutive days of spontaneous differentiation: the well that was seeded first had been undergoing the differentiation protocol for 7 days, and for the well that was seeded last the differentiation protocol was never initiated, making it undergo 0 days of differentiation. Next, cells were incubated with Hoechst (1:1000), NL493-conjugated Mouse Anti-Human TRA-1-81, NL557-conjugated Mouse Anti-Human SSEA-1 and Alexa Fluor 647-conjugated Mouse Anti-Human NCAM-1/CD56 live stains diluted in media (1:50) for 30 minutes. Confocal images of live colonies were acquired on a Nikon UltraVIEW VoX W1 Spinning Disk Confocal with sCMOS camera at 10x magnification (0.65 mm/px), 100 ms exposure and 100% laser power for all wavelengths. Next, cell culture media and silicone well were removed, and samples were washed by submerging the plate into 5 mM ammonium formate buffer for 3s to enhance spectral abundances. Norharmane was used as MALDI matrix and deposited via sublimation. A slide containing cell colonies was taped to the bottom of the condenser in a simple sublimation apparatus. Solid norharmane was placed at the bottom of such sublimation apparatus. Sublimation was performed at 250 C under vacuum for 6 min.

### S6. MALDI TOF MS imaging

Matrix-deposited samples were analyzed in reflectron mode using a RapifleX Tissuetyper time of-flight (TOF) mass spectrometer (Bruker Daltonics, Billerica, MA, USA) equipped with a Smartbeam3D 10 kHz Nd:YAG (355 nm) laser. Imaging experiments were controlled by the FlexImaging 4.0 software (Bruker Daltonics, Billerica, MA, USA) using the single smartbeam laser setting (∼5 μm in both x and y dimensions) with a laser raster size of 10 μm in both x and y dimensions. Data were collected in negative ion mode in the *m/z* 200-1600 range with 200 laser shots averaged at each pixel. Mass calibration was performed using red phosphorus as a standard prior to data acquisition.

### S7. MALDI FTICR MS experiments

Ultra-high mass resolution data was collected on a Bruker solariX 12-Tesla Fourier-transform ion cyclotron resonance (FTICR) mass spectrometer equipped with a MALDI ion source. Data was acquired in negative mode from *m/z* 300-1200 at 1M transient size with 25 µm raster width. Laser was set to minimum focus at 25% power. Real time calibration was employed with lock masses 333.11457 (deprotonated norharmane dimer) and 885.54986 (deprotonated PI 38:4). Data pre-processing was done in SCiLS Lab (SCiLS GmbH, Bremen, Germany) software. Mass spectra were preprocessed during import into SCiLS Lab by converting to spectra to centroid. MS/MS data was collected using quadrupole precursor mass selection. Collision energies ranged from 15-35 eV for selected peaks.

### S8. Co-registration and segmentation

All MALDI-MS data preprocessing was performed using the SCiLS Lab (SCiLS GmbH, Bremen, Germany) software. Mass spectra were preprocessed during import into SCiLS Lab using baseline removal by iterative convolution. A minimum interval width of 20 mDa around the average peak center was used to account for peak shifts throughout the experiment. Automated pick peaking was first performed using SCiLS Lab, followed with manually peak screening to select the *m/z* features that were associated with the cell colony distribution. A custom Python script using *imzmlparser* library (https://github.com/alexandrovteam/pyimzML) was used to extract *m/z* spectra for each pixel from imzML and .bd files generated by the RapifleX instrument. Each peak at *m/z* value of interest was integrated at ±0.4 Da to receive the final ion abundance value. Next, we used the co-registration GUI created in our lab to align confocal and MALDI imaging data and extract cell by cell *m/z* spectra. We used a confocal image stained with Hoechst nuclei live dye and a MALDI ion image averaged over the *m/z* spectrum as reference images for alignment. The algorithm rotates, shifts, and scales reference images in a given range of parameters until the global maximum of mutual information of the images is found. The confocal image stained with Hoechst was also used to segment nuclei to extract the abundance data at each *m/z* value of interest on a cell-by-cell basis along with the corresponding fluorescence intensities from the confocal image. First, a local threshold was applied on the image at window sizes ranging from a fourth of the image to twice the size of the largest cell (parameter provided by user); this was done to obtain the most comprehensive binary mask of the nuclei. Next, the segmentation algorithm utilizes a multiscale Laplacian of Gaussian (LoG) blob detection algorithm [21] implemented using OpenCV [22] for Python to find nuclei seeding points. The LoG of the image is computed for each radius in a user-provided range of nuclear radii. The LoG is found by applying a Gaussian filter with a standard deviation of 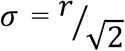, where r is the radius in pixels, and subsequently finding the second spatial derivative of the image. This results in a range of images containing several local minima at the center of each blob; the intensity of the minimum at each blob corresponds with how closely the actual radius of the blob matches the *σ* parameter of the Gaussian filter. Following normalization of each image by multiplying it by *σ*^2^, the minimum at each pixel across the stack of all calculated LoGs is taken. The center of each nucleus, or seed, can be found at the local minima of this resultant image, and applying the watershed transformation at these points on the binary mask yields the nuclear labels.

### S9. Metrics

#### Cell-to-cell connection distance

is calculated by first finding a closest neighbor distance for each cell. Next, we clean these distances from outliers – remove all values that are higher than mean plus three standard deviations. Finally, the cell-to-cell connection distance value is assigned the maximum of the cleaned distances array. This approach guarantees that every non-outlier cell will have at least one cell within the cell-to-cell connection radius. The cells within that radius are called neighbors.

#### Neighbor-relative abundance

was measured by first finding all neighbors for a cell of interest. Next, average abundance is calculated among the neighboring cells, and the self-value is divided by the average neighbor value.

To identify *dividing cells*, we segmented the nuclei images and calculated the following metrics for each cell: neighbor-relative Hoechst intensity, the area of the nucleus, and the distance to the nearest neighbor. A K-Means (K = 2) clustering algorithm from the scikit-learn library was then trained to classify the cells as either ‘dividing’ or ‘not dividing’ using those metrics. Out of the two resulting class centers, we designated a center with higher neighbor-relative Hoechst intensity, smaller area, and larger nearest neighbor distance to represent the dividing cells class. To estimate the classification accuracy, we manually annotated 3 1000×1000-pixel patches of Hoechst-stained colony image (Fig. 6 – supp. 2) and used the algorithm to predict cell labels, yielding an accuracy of 98.8%. Sensitivity of 72.5% and a positive predictive value of 96.3% showed that this method is much more prone to false negatives than false positives, which is preferrable when data has a disproportionately high number of negative datapoints.

To detect the *edge of the colony* we multiplied cell-to-cell connection distance by the user-provided value (default is 3) to expand the cell’s neighborhood. Cells that are on the edge of a colony can be distinguished by having at least one side with no neighbors in its network. To determine if the cell is on the edge, a cell’s personal neighborhood is represented as a series of vectors connecting the center node cell and each of its neighbors. Next, we sort these vectors by their angles, and if any difference between two consecutive angles is greater than π/2 radians, the cell is labelled as an “edge” cell. Next, the *edge distance* metric can be derived by finding the distance between a given cell and the edge cells and taking the minimum value.

### S10. Classification tree

Out of each day of differentiation we randomly selected 8000 cells and merged it into a training dataset of 64000 data points, with differentiation day number as a label and *m/z* values as features. We repeated this with a replica experiment obtaining a validation dataset of 64000 data points. Classification tree for the day of differentiation prediction was built using MATLAB built-in function *fitctree* with the number of tree splits limited by *MaxNumSplits* parameter set to 20. We used variable appending method for variable selection, iteratively adding those variables that increased the prediction accuracy on a validation dataset the most after being included into the analysis. After introducing first 5 variables this way the accuracy has stopped increasing, this indicated that these variables are the most predictive of the differentiation time point.

### S11. Partial least squares discriminant analysis

To determine which cells are TRA-1-81, SSEA-1 or NCAM-1 positive we first applied k-means clustering analysis to the corresponding extracted fluorescence intensities using MATLAB built-in function *kmeans* with number of clusters equal 2 for each live stain individually. NCAM-1 signal was spatially clustered and had high contrast between positive and negative population, thus all NCAM-1 positive cells were marked as negative in other stains for the purpose of PLS-DA. All NCAM-1 negative cells were assigned 2 different labels: positive or negative in TRA-1-81 and positive or negative in SSEA-1. Next, we excluded double negative and double positive cells from the analysis as potential artifacts of staining and assigned all the remaining cells a pluripotent versus differentiated label, where pluripotent cells are positive in TRA-1-81 and negative in SSEA-1 and vice versa for the differentiated cells. After all cells received their lineage label, we used SIMCA® software (Sartorius AG, Göttingen, Germany) for PLS-DA to predict the labels using *m/z* values as features. To select the best predictors, we used variable trimming: iteratively removing every variable that reduced prediction accuracy on a validation dataset.

### S12. Image processing

All microscopy images were acquired with NIS-Elements Viewer software (Nikon Instruments Inc., Melville, NY, USA). Channels were independently exported as grayscale TIFF files, then uploaded to ImageJ [23], merged with consistent contrast settings for each antibody, and saved as RGB JPEG files. Data processing for MS image creation is described in Methods S8. MS images were saved by OpenCV’s *imwrite* command as grayscale PNG files, then uploaded to ImageJ with contrast adjusted consistently for every *m/z* value and saved using “jet” colormap. Both microscopy and MS images were cropped to the overlapping region after alignment.

**Figure 6 – figure supplement 1.**
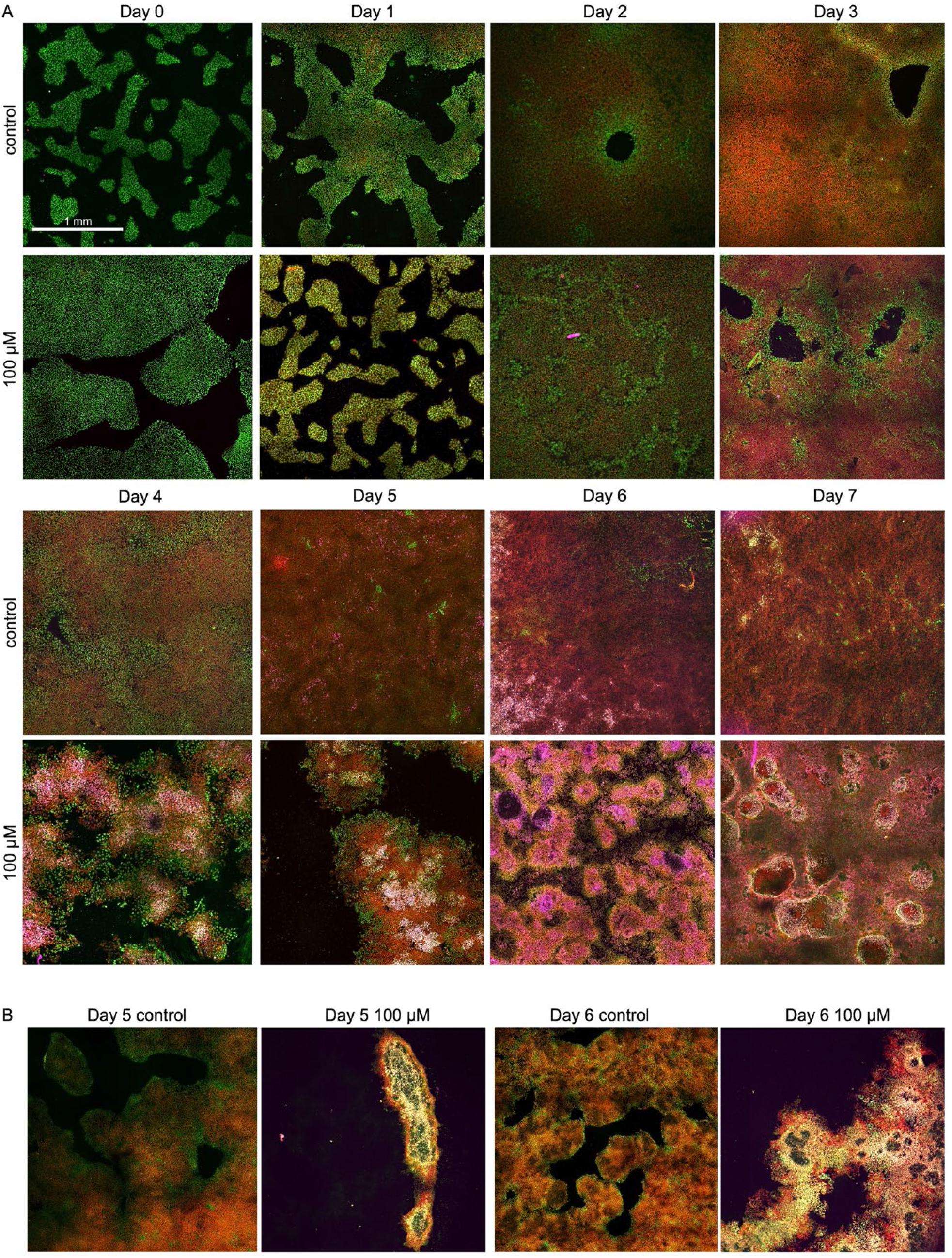
Immunocytochemistry images of the 8-day spontaneous differentiation for control and PI 3-kinase inhibited conditions. Cells were fixed and stained for Oct4 (green), Otx2 (red), and Pax6 (pink) in an independent set of experiments.

**Figure 6 – figure supplement 2.**
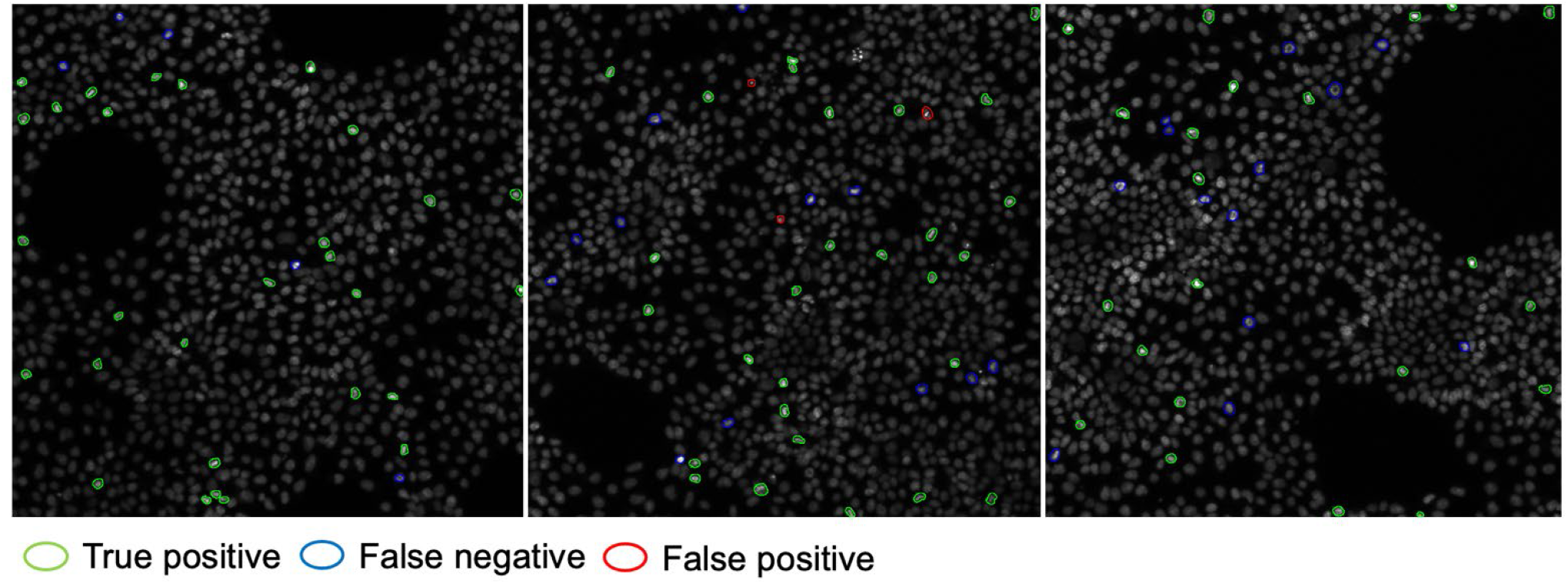
Dividing cells manual annotation and validation results. Circled nuclei were manually annotated as dividing (109 cells total). Segmentation algorithm detected a total of 2846 cells and neighbor-relative Hoechst intensity, area, and nearest neighbor distance were calculated for each cell. Using those metrics, K-means clustering predicted the dividing cells with accuracy of 98.8%, sensitivity of 72.5% and a positive predictive value of 96.3%.

**Figure 7 – figure supplement 1.**
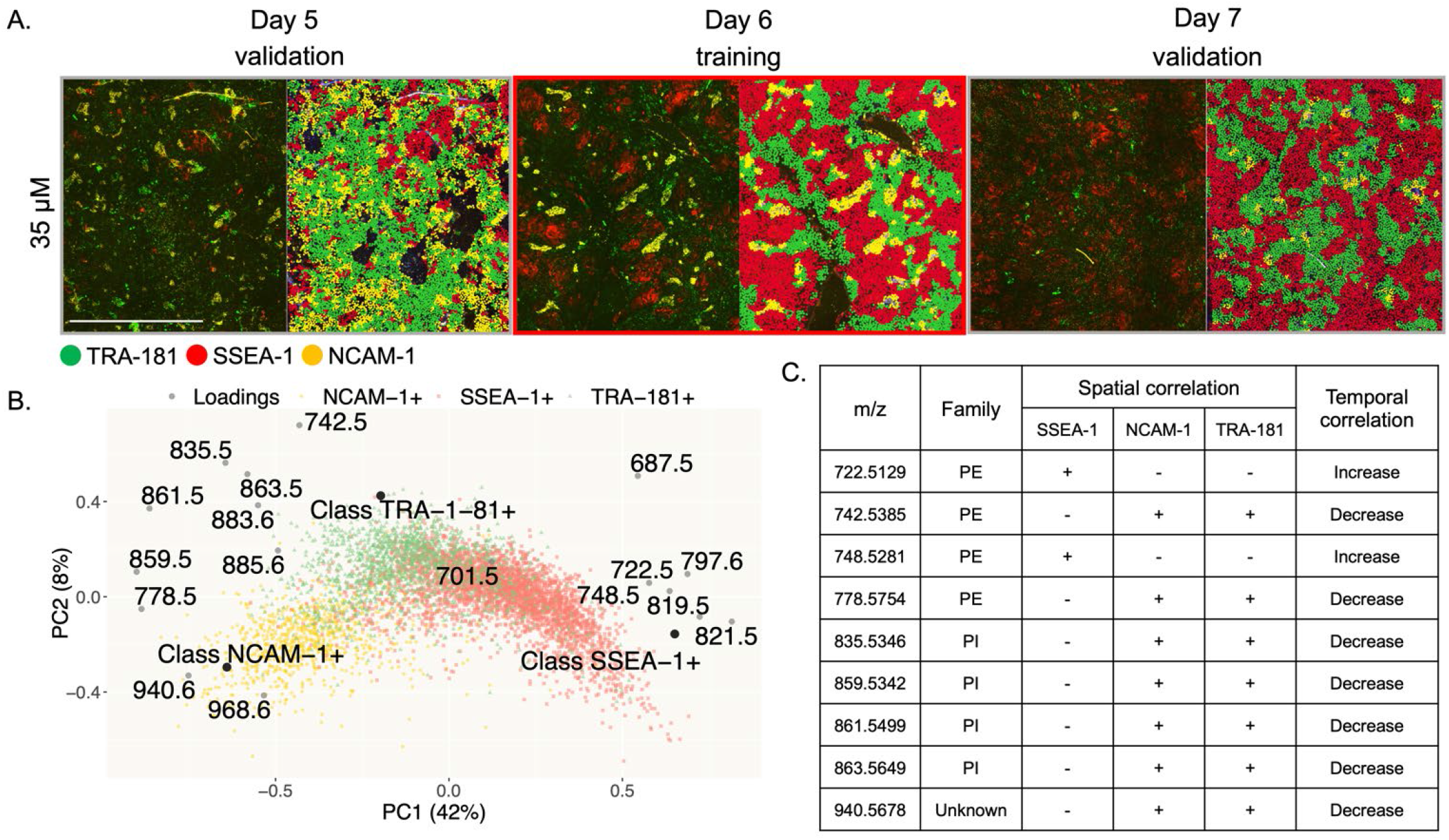
Confocal images of days 5-7 of spontaneous differentiation with 35 µM of LY294002 and their predicted lineage labels. Green color labels TRA-181+ cells, red color labels SSEA-1+ cells, yellow colors NCAM-1+ cells. Day 6 was used as a training set, days 5 and 7 as validation sets (66% and 71% accuracies). Scalebar is 1 mm. B. Biplot of the PLS-DA model used to discriminate between the cell populations above. C. Summary table of lipid ions consistently correlating with cell fate. See Table 2 for expanded annotation.

**Figure 6 – Source data 1.**
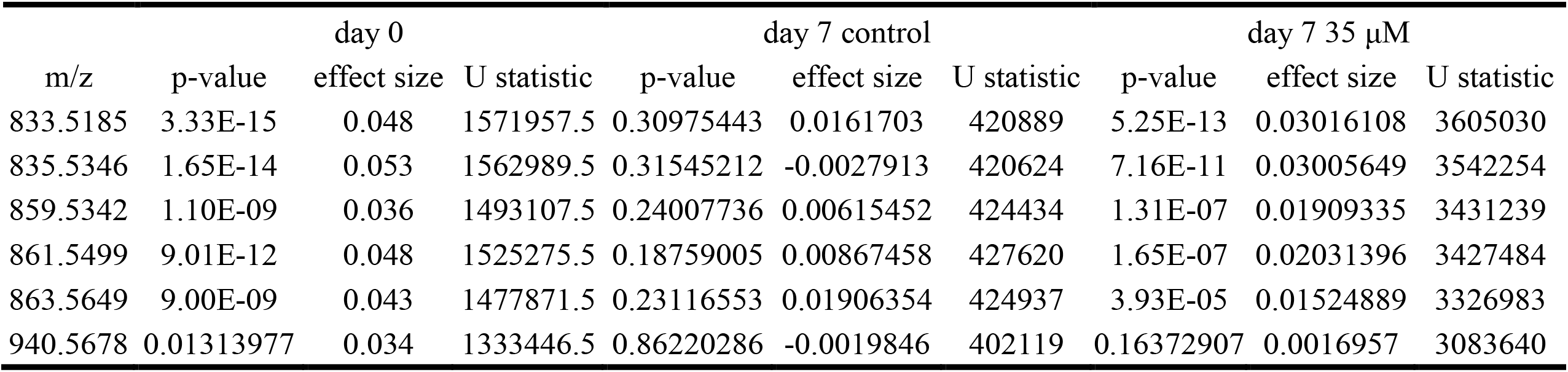
Due to non-normal distribution of data Mann-Whitney U test was used for comparison of neighbor-relative abundance of phospholipids in dividing versus non-dividing cells. The table summarizes resulting statistics. Effect sizes are calculated as a difference between median value for dividing cells and median value for non-dividing cells.

